# High-resolution MEMRI characterizes laminar specific ascending and descending spinal cord pathways in rats

**DOI:** 10.1101/781823

**Authors:** Vijai Krishnan, Jiadi Xu, German Alberto Mendoza, Alan Koretsky, Stasia A Anderson, Galit Pelled

## Abstract

Manganese Enhanced MRI (MEMRI) utilizing different manganese chloride (MnCl_2_) delivery methods, has yielded valuable architectural, functional and connection information about the brain. MEMRI also has the potential in characterizing neural pathways in the spinal cord. The spinal cord grey matter is anatomically composed of nine distinct cellular laminae, where each of the laminae receives input from a specific type of neuronal population and process or serves as a relay region in a specific sensory or motor pathway. This type of laminar arrangement in the spinal cord is currently only visualized by histological methods. It is of significant interest to determine whether laminar specific enhancement by Mn^2+^ can be achieved in the spinal cord, as has been reported in the brain and olfactory pathway. Here we focus on using MEMRI to determine the specific laminae of the thoracic region of the spinal cord. We focus on MnCl_2_ changes in the ascending and descending tracts of the spinal cord. Major factors in applying this technique in the spinal cord are the ability to acquire high-resolution spinal cord images and to determine a noninvasive route of administration which will result in uptake by the central nervous system.

We have applied the MEMRI approach by intraperitoneal (i.p). delivery of MnCl_2_ and imaged lumbar and thoracic spinal cord levels in rats to determine whether T_1_ weighted MRI can detect spinal cord laminae 48 hours following MnCl_2_ administration. T_1_ weighted images of the lower lumbar level were obtained from MnCl_2_ injected and control rats. Here we demonstrate laminar specific signal enhancement in the spinal cord of rats administered with MnCl_2_ vs. controls in MRI of the cord with ultra-high, 69 μm in-plane resolution. We also report reduced T_1_ values over time in MnCl_2_ groups across laminae I-IX. The regions with the largest T_1_ enhancements were observed to correspond to laminae that contain either high cell density or large motor neurons, making MEMRI an excellent tool for studying spinal cord architecture, physiology and function in different animal models.

## Introduction

The spinal cord consists of nine laminae that have different cytological and functional characteristics. The large somatic motor neurons that occupy laminae IX are essential for maintaining muscle tone and innervating muscle spindles (1). Neuronal death in that region can lead to a number of motor neurons disorders, such as amyotrophic lateral sclerosis (ALS) (2, 3), spinal muscular atrophy, spinal bulbar muscular atrophy and spinal muscular atrophy with respiratory distress 1 (4). To date, motor neuron diseases are untreatable and may be fatal, as in ALS, or cause severe clinical symptoms. Most of the motor neuron diseases initially affect only a specific cell population within the spinal cord and does not result in gross architectural changes. Using non-invasive imaging tools to monitor the viability of the large motor neurons in the spinal cord could greatly contribute to the ongoing efforts of finding new therapeutic strategies for motor neuron disorders. There is an ongoing need to develop technologies that will allow non-invasive imaging of spinal cord function. Nevertheless, in *vivo* high-resolution imaging of the spinal cord remains a challenge due to several factors such as the susceptibility to motion, variation in tissue depth over the length of the cord, and artifacts from the cerebrospinal fluid (CSF) and vertebrae.

Manganese enhanced magnetic resonance imaging (MEMRI) has been applied to identify and trace connectivity changes in specific neuronal architecture of the brain and spinal cord. MEMRI has also used to assess functional information pertaining to different areas of the brain affected by a variety of brain diseases (5-8). The Mn^2+^ uptake by neurons has been shown to be tightly coupled to the neuronal activity (9-14), and hence, the cells viability. Mn^2+^ ions have been shown to accumulate in the desired tissue leading to a subsequent shortening of T_1_ weighted intensity. This in turn leads to a favorable contrast enhancement in T_1_-weighted MRI signaling in the tissue. Therefore, changes in T_1_-weighted signal values of different spinal cord regions after Mn^2+^ delivery may provide an indication of the cells function. Indeed, it has been demonstrated that MEMRI can detect lesions and Mn^2+^ transport within spinal cord neurons can give an indication of the degree of the spinal cord recovery in rodents (15-17).

Here we demonstrate that 48 h following systemic injections of MnCl_2_ in rats, there is a significant accumulation of Mn^2+^ ions within localized spinal cord laminae, specifically in laminae that consist of large motor neurons, as well as in laminae that consist of particular high cell density. We also report that MEMRI enhanced visualization of laminae regions I-IX of the thoracic segment of the rat spinal cord. These findings were further correlated with immunostaining. These findings can be used to assess discrete laminar specific changes of the spinal cord *in vivo*.

## Methods

### MnCl_2_ administration

Animal procedures were performed in accordance with Johns Hopkins animal care and use committee guidelines. Male Sprague-Dawley rats (200g) were anesthetized with 2% Isoflurane and 2 ml of 100 mM MnCl_2_ dissolved in saline was delivered i.p. with 0.4 mmol/kg of body weight (n=8). Control rats received only saline injections (n=3).

### Animal preparation for MR

48 hours following MnCl_2_ administration, rats were anesthetized with 2% Isoflurane and were placed supine (in order to suppress motion) in a secured MR head and body cradle. A 4×1 phased receiving coil was placed under the rats thoracolumbar vertebrae. The head was slightly stretched and tilted back to reduce curvature at the cervical region. The head, body, and legs were secured with tape during the experiment and the breathing rate was monitored. Each rat was imaged before and after MnCl_2_ administration.

### MRI

An ultra-high field 11.7 Tesla/16 cm horizontal bore small-animal scanner (Bruker BioSpin, Rheinstetten, Germany) was used for imaging. A 72 mm quadrature volume resonator was used as a transmitter. T_1_ weighted images were collected using rapid acquisition with refocused echoes (RARE) imaging module with an echo time of 14 ms, RARE sequence with 2 echoes, slice thickness=1 mm, a matrix size of 320 × 320 and a field-of-view (FOV) of 2.2 × 2.2 cm. T_1_ maps were measured using a RAREVTR sequence (RARE with six variable repetition times 500, 700, 1000, 1500, 2000, 5000 ms). The matrix size and FOV were identical to the T_1_ weighted images. 17 slices were acquired, beginning at the lower lumbar area and moving up towards the cervical region. Three saturation slices around the spinal cord area were used to allow for a reduced FOV around the cord without alias. A fat suppression module was utilized in all experiments.

### Histology

At the end of the imaging session, rats were perfused with 200 ml of saline and then 200 ml of 4% paraformaldehyde in 0.1M phosphate buffer (PBS), pH 7.4. The lumbar area of the spinal cord was removed and post-fixed in perfusion solution for 2 h at 4°C and cryoprotected in 30% sucrose solution in PBS. Sectioning was done with a freezing microtome (50 µm thick slices) and sections were mounted onto slides. Sections were thawed to room temperature and blocking (3×10 min) was done with PBSG-T (0.1M PB pH 7.4 + 0.9% NaCl + 0.2% gelatin + 0.2% triton). Neuronal nuclei were incubated with NeuN monoclonal antibody (Chemicon) at a 1:1000 dilution for 2 h at room temperature. Primary antibody was incubated overnight at 4°C and then washed 3 × 20 min in PBS, pH 7.2. Immunoreactivity was visualized with the appropriate secondary antibody conjugated to Alexa 488 (1:1000 dilution; Molecular Probes, Eugene, OR). Sections were counterstained with DAPI, which labels all cell nuclei. Sections were rinsed (3×10 min) in PBSG-T and incubated with an anti-mouse biotinylated secondary antibody (1:500, Chemicon) for 45 minutes at room temperature. For DAB detection, sections were rinsed (3×10min) in PBSG-T and incubated with Strepavidin HRP diluted in Tris buffered saline (1:500, Chemicon) for 30 minutes at room temperature. Sections were rinsed (2×10 min) with phosphate buffered saline + 0.2% triton, and (2×10 min) with phosphate buffered saline alone. Detection was performed with DAB chromagen reagent (Chemicon) incubated for 1-5 minutes and rinsed with PBS when staining was optimal. All images were acquired at 20× magnification using a computer-controlled microscope (Zeiss Axio Observer Z.1).

### Data Analysis

The mean T_1_ values and standard error was calculated for each region-of-interest (ROI). T_1_ values were calculated using Matlab. T_1_ values are represented as mean ± SEM. Data was analyzed by paired t-test and results were considered significant if p<0.05.

## Results

Systemic administration of MnCl_2_ through i.p injections caused efficient uptake of Mn^2+^; our results show that MnCl_2_ administration provides visual enhancement of the different regions of the spinal cord. Analysis of MRI-T_1_ maps show significant changes post MnCl_2_ administration. These shorter T_1_ values serves as the basis for contrast enhancement and delineating specific anatomical structures in the spinal cord.

**Figure 1** demonstrates T_1_ weighted images of rat spinal cord before and after MnCl_2_ injection. Images are from L4 lumbar vertebrae level obtained with 69 µm in-plane resolution. T_1_ weighted images of spinal cord from another Mn^2+^ injected rat is shown in **Figure 1B** with the corresponding coronal anatomical section. Interestingly, the superficial layers, which consist mostly of the terminations of primary afferent nociceptive fibers and neurons of lamina I and substantia gelationsa can be visualized in the control MRI slice but are much more enhanced in Mn^2+^ enhanced MRI slice. In rats that received Mn^2+^, additional T_1_ enhancement can be seen in the ventro-lateral grey matter, which is a region that contains large motor neurons.

**Figure 1.**
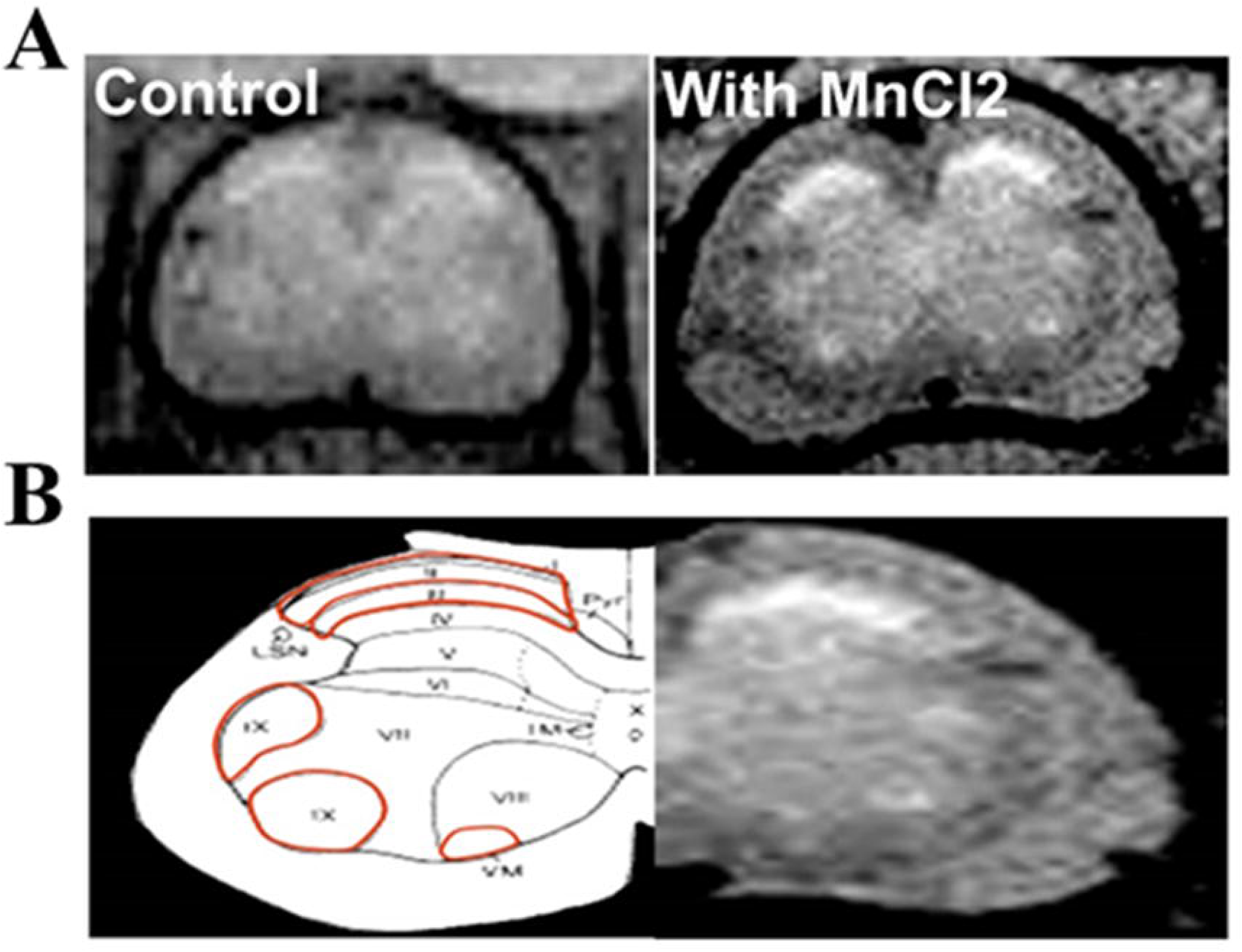
**(A)** T_1_ weighted images of coronal L4 spinal level in control and Mn^2+^ injected rats acquired 48 hours following Mn ^2+^ i.p. administration showing increased contrast enhancement. **(B)** Shows half of the cord from a Mn^2+^ injected rat labelled with the corresponding laminae anatomical sections.

### Reduction of T_1_-Signals in MnCl_2_ injected rats show increased contrast enhancement of grey matter

MRI images of thoracic segments were obtained 48 h after MnCl_2_ injection. Signal enhancement in different spinal cord regions was calculated and normalized according to the nearby muscle. Signal enhancement within grey matter layers and proximate grey matter regions that contain small diameter interneuron populations in Mn^2+^ injected rats and control are demonstrated in Figure 2. **Figure 2A** shows three thoracic segments and anatomically defined ROIs. Laminar regions (I-III), (V-VII) and (VIII-IX) correspond to ROIs (3,6,9); (2,5,8) and (1,4,7) respectively. **Figure 2B** show correlation fit curves for signal intensity values at six different repetition times for all measured ROIs. **Figure 2C** shows that the mean T_1_ values were significantly different before (1.5627 ± 0.01s) and after MnCl_2_ (1.4411 ± 0.02s) administration (p=0.0007) in laminae I-III region. In Laminae V-VII region, there was a significant decrease in T_1_ (pre-MnCl_2_: 1.5508 ± 0.017s; post-MnCl_2_: 1.43511 ± 0.025s) (p=0.02) as well. T_1_ values in VIII-IX region were 1.56 ± 0.02s before MnCl_2_ administration. Post MnCl_2_, T_1_ values were reduced to 1.442 ± 0.021s, p=0.0007.

**Figure 2.**
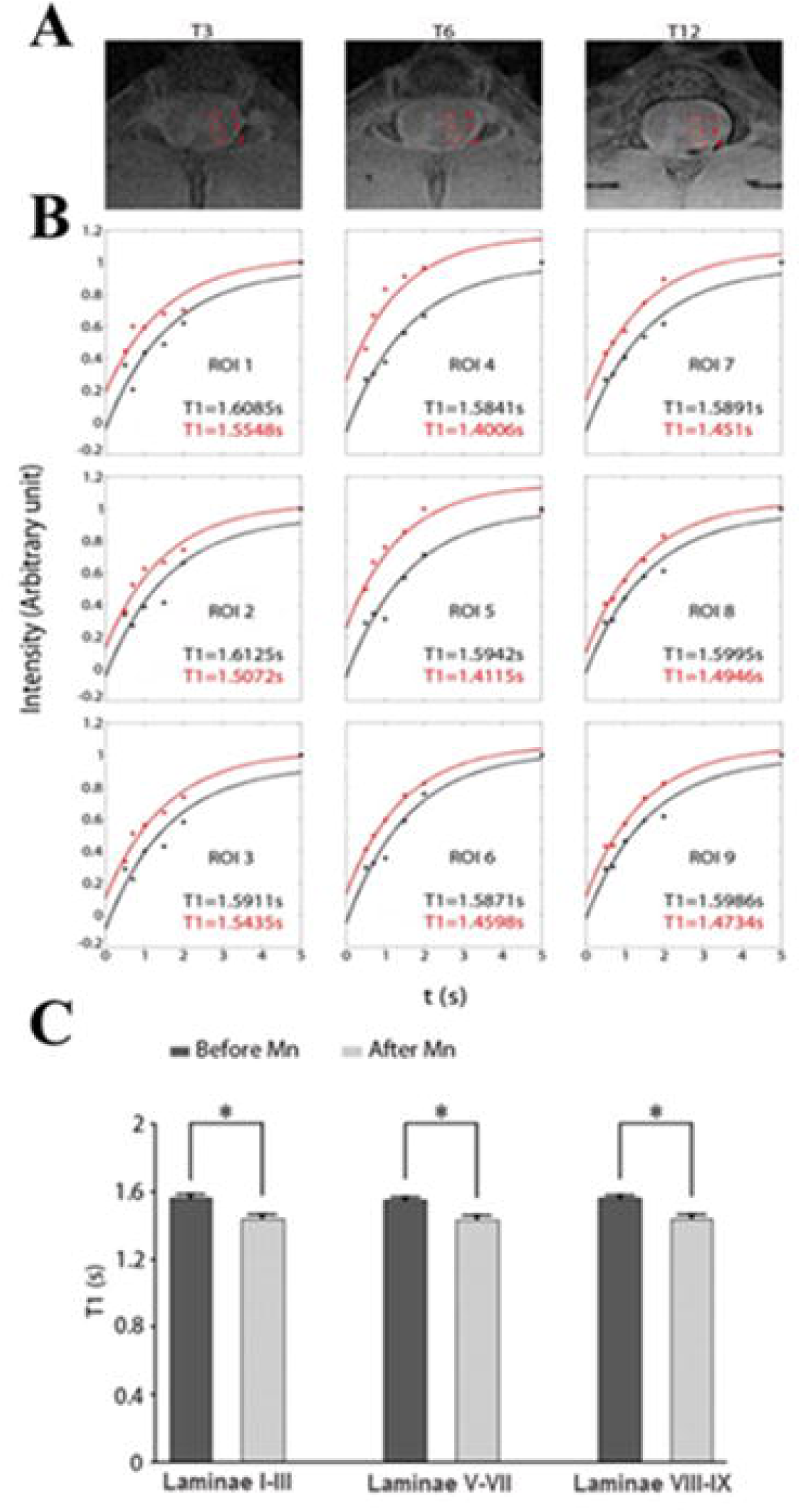
**(A)** Specific laminae regions across the thoracic section of the rat spine showing regions of interest (ROI) 1-9 (**B)** Individual representation of T_1_ values across ROI regions 1-9 before (black) and after (red) MnCl_2_ administration, with post-MnCl_2_ groups showing lower T_1_ values **(C)** Grouped pre-MnCl_2_ laminae T_1_ average compared to post-MnCl_2_ laminae average. Paired T-Test analysis of laminae were significantly different (p<0.05).

### Reduction of T_1_ signal in corticospinal and spinalcortical white matter post MnCl_2_ administration

We sought to calculate the signal enhancement in white matter tracts post MnCl_2_, specifically in regions of ascending and descending tracts. **Figure 3A** shows a representative MnCl_2_ image with ROIs corresponding to spinalcortical (sensory pathway that conveys impulses concerning position of different body parts) and corticospinal (involved with voluntary motor function and sensory impulses) regions. Data from **Figure 3B** shows mean T_1_ values for spinalcortical and corticospinal white matter regions before and post MnCl_2_ treatment. The changes in the mean T_1_ values for the spinalcortical region (Red ROI in middle) before (1.5458 ± 0.016 s) and after (1.4629 ± 0.025 s) MnCl_2_ were significant (p=0.003). The descending motor pathway (corticospinal) also had a significant visual enhancement (pre-MnCl_2_, 1.5368 ± 0.02 s) vs (post-MnCl_2_, 1.4467 ± 0.02 s), p=0.002. Thus, our results show that MEMRI is effective in the contrast enhancement of regions that pertain to both sensory and motor pathways.

**Figure 3.**
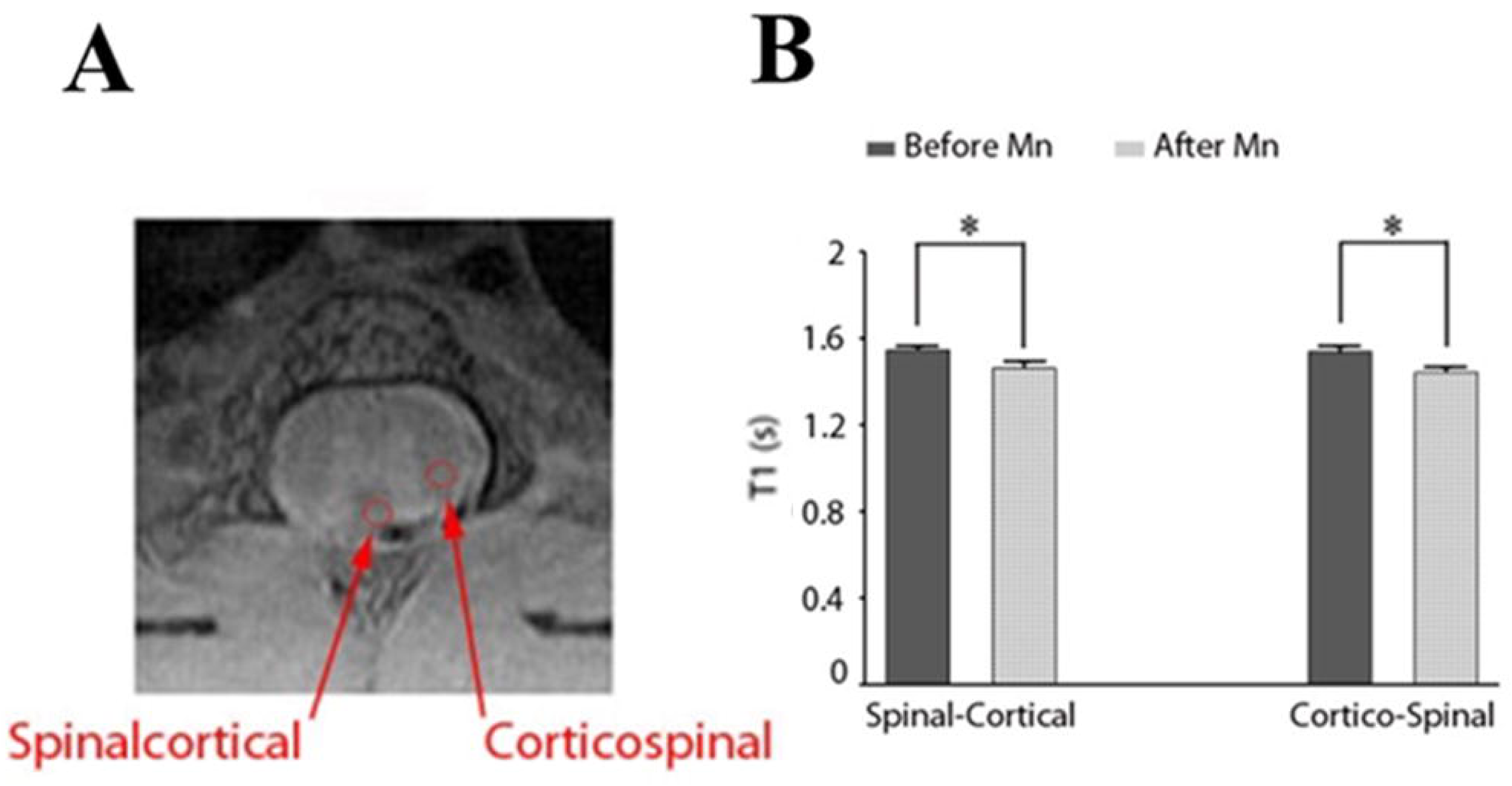
**(A)** Specific regions (red ROI) of the spinal cord that correlate to ascending and descending tracts. **(B)** Grouped pre-MnCl_2_ white matter averaged areas compared to post-MnCl2 averaged white matter areas. Paired T-Test analysis of white matter tracts, pre and post, were significantly different (p<0.05).

Next, we used histological methods to delineate specific regions of interest in the spinal cord slices obtained from the same rats following the imaging session. DAB staining of coronal MnCl_2_ administered L4 T_1_ regions revealed areas of high cell density **(Figure 4A)**. This was reflected as a hyper-intense MRI signal (area outlined in black). The second high intensity MRI region was identified as an area that has a large population of motor neurons (area outlined in red). Both histology and corresponding MRI images are placed side by side for comparison. **Figure 4B** further delineated in detail the regions identified in **Figure 4A.** The boxed region labelled A was identified as laminae regions I × II, which contain a high density of cells that responds to noxious and thermal stimuli (18). **Figure 4A** also shows laminae IX region with the presence of large neurons. These have previously been identified as somatic motor neurons that innervate muscles (19). **Figure 4C** shows effective labelling of the spinal cord with mature neuronal marker (NeuN) and nuclear staining (DAPI). Extensive labelling with NeuN was seen in both dorsal and ventral grey matter of the spinal cord. Double labelling with NeuN and DAPI were visualized and reported.

**Figure 4.**
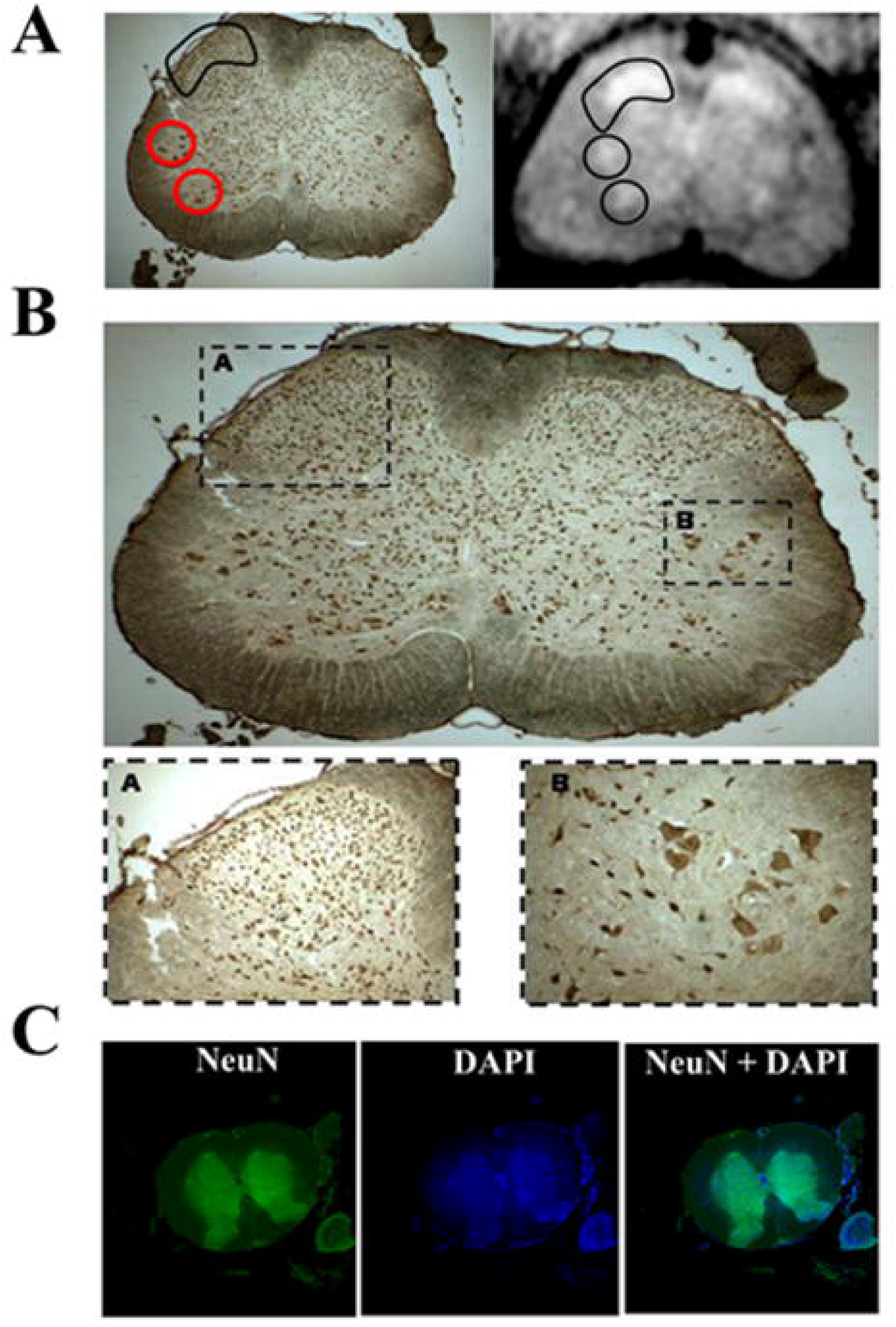
**(A)** Coronal L4 T_1_ weighted image with the corresponding histology taken from a Mn^2+^ injected rat. Note that the hyper-intense MRI regions match regions where high cell density (black) and large motor neurons are found (red). **(B)** A coronal L4 histology image. Laminae I & II have high density cell bodies (box labelled A) and respond strongly to noxious and thermal stimuli. Lamina IX (box labelled B) consists of several distinct clusters of large somatic motor neurons that innervate muscles in the lumber level. Boxed sections A and B has been enlarged to show hyper intense cell bodies and motor neurons. **(C)** Coronal sections showing neuronal labelling (NeuN in green), nuclei labelling (DAPI in green) and merged image of NeuN and DAPI.

## Discussion

The use of MEMRI for in vivo neuronal tracing has been on the rise. Here, we showed that MEMRI can visually enhance specific regions of the spinal cord through decreased T_1_ changes post MnCl_2_ administration. Our results demonstrate efficient Mn^2+^ transport and accumulation in corticospinal and spinalcortical pathways after i.p. injection.

Different spinal cord regions are associated with various neurological disorders such as ALS, multiple sclerosis, nerve damage and chronic pain. There is a growing interest in using cellular and stem cell therapies to combat different motor neurons disorders (20). Currently, the only approaches to assessing the efficacy of these strategies has either been behavioral testing or histology with *ex vivo* imaging methods. The possibility of imaging *in vivo* and non-invasively the function of different cell population within the spinal cord grey matter, opens a new frontier in the assessment of different therapeutic strategies.

Pathologies of the spinal cord tracts have been detected using diffusion tensor and functional magnetic imaging techniques (21, 22) and MEMRI (15, 16). Recent reports have focused on using MEMRI to assess neuronal damage post spinal cord injury (23) and in assessing signal transduction mechanisms in the spinal cord (24). These reports demonstrate the applicability of the techniques to follow gross architectural and to some degree, functional changes of the spinal cord.

It has been shown that Mn^2+^ ion enter cells through voltage gated calcium channels (10, 25). Different classes of these channels are used to amplify and increase the output of the cells. At the lumber 4 level of the spinal cord, the motor neurons in lamina IX are larger in size compared to other neurons in the spinal cord grey matter at that level. Since their surface area is large (more than 500,000 µm^2^) they exhibit more Ca^2+^ channels per cell and therefore the accumulation of Mn^2+^ within them is greater. In contrast, lamina I is composed of small and medium sized cells that respond specifically to noxious stimuli, and lamina II of the spinal cord is composed of tightly packed small cells that are also involved in pain processing. However, dense cell architecture in these laminae may underlie the significant enhancement in the signal of that region. Interestingly, there is an enormous interest both in visualizing cells function in the upper dorsal horn laminae for drug development that targets acute and chronic pain, as well as in the large motorneurons in lamina IX in respect to following therapeutic strategies for motorneurons disorders.

Systemic manganese infusion was applied previously to study cerebral architecture in normal (26) and specific mutation in rodents (27, 28). Indeed, a difference in the cerebral architectural of the mutant mice was visualized using this method. We show that MEMRI is successful in diffrentiating different grey matter regions within the spinal cord and can be used for longitudinal studies of spinal cord physiology *in vivo*.

## Author contributions

VK, JX, GAM, AK, SA and GP designed experiments, acquired and analyzed data. VK and GP wrote and the edited the paper.

## Acknowledgements

This work was supported by National Institute of Health/ National Institute of Neurological Disorders and Stroke R01NS098231 and NIH/NINDS R01NS072171 grants. The authors thank the Intramural National Institute of Neurological Disorders and Stroke and the National Heart, Lung, and Blood Institute for support

